## Supplementary material for "E.PathDash, pathway activation analysis of publicly available pathogen gene expression data": Supplmental Figures S1-S10, Table S1

### E.PathDash, pathway activation analysis of public cystic fibrosis pathogen gene expression data

*Supplementary Information*

---

| Name | File Type | Contents | Access Point |
|---|---|---|---|
| Raw gene count data | zip | Matrix of gene counts for each dataset, design matrix for study samples | Study Explorer |
| Differential gene expression data | csv | Results of differential gene expression analysis for selected dataset and treatment comparison (gene, logFC, p-value) | Study Explorer |
| Significant KEGG pathways box plot | png | Box plot of distributions of gene logFC values for significantly expressed KEGG pathways for selected datasets and treatment comparison | Study Explorer |
| Significant GO terms box plot | png | Box plot of distributions of gene logFC values for significantly expressed GO terms for selected dataset and treatment comparison | Study Explorer |
| KEGG pathway analysis table | csv | Table of all KEGG pathways analyzed in pathway analysis (binomial test statistic, median gene logFC, p-value, FDR corrected p-value) | Study Explorer |
| GO term analysis table | csv | Table of all GO terms analyzed in pathway analysis (binomial test statistic, median gene logFC, p-value, FDR corrected p-value) | Study Explorer |
| KEGG pathway datasets table | csv | Table of all datasets and treatment comparisons with significant expression of selected KEGG pathway (study identifier, treatment comparison, median gene logFC) | KEGG Pathway Explorer |
| KEGG pathway volcano plot | png | Plot of logFC value and log-transformed p-value for genes along selected KEGG pathway in selected dataset and treatment comparison | KEGG Pathway Explorer |
| Differential expression of KEGG pathway genes | csv | LogFC and p-values for genes along selected KEGG pathway | KEGG Pathway Explorer |
| GO term datasets table | csv | Table of all datasets and treatment comparisons with significant expression of selected GO term (study identifier, treatment comparison, median gene logFC) | GO Term Explorer |
| GO term volcano plot | png | Plot of logFC value and log-transformed p-value for genes along selected GO term in selected dataset and treatment comparison | GO Term Explorer |
| Differential expression of GO term genes | csv | LogFC and p-values for genes along selected GO term | GO Term Explorer |
| Study comparison bar chart | png | Bar plot of median gene logFC values for each sample comparison within selected study for selected KEGG pathway or GO term | Study Comparison |

**Table S1.**

Downloadable content available in E.PathDash. “Access Point” refers to the page of the application where the content can be accessed.

**Table S2.**

LogFC and p-values for genes in the propanoate metabolism KEGG pathway for *P. aeruginosa* grown in co-culture with WT *C. albicans* compared to monoculture and for *P. aeruginosa* grown with *adh1*Δ/Δ *C. albicans* compared to WT *C. albicans*. Gene and protein names as well as other annotation information retrieved from uniport.org based on Uniport IDs.

*Table S2 is available in a separate supplementary file titled "Supplemental_Table_2.xlsx"*

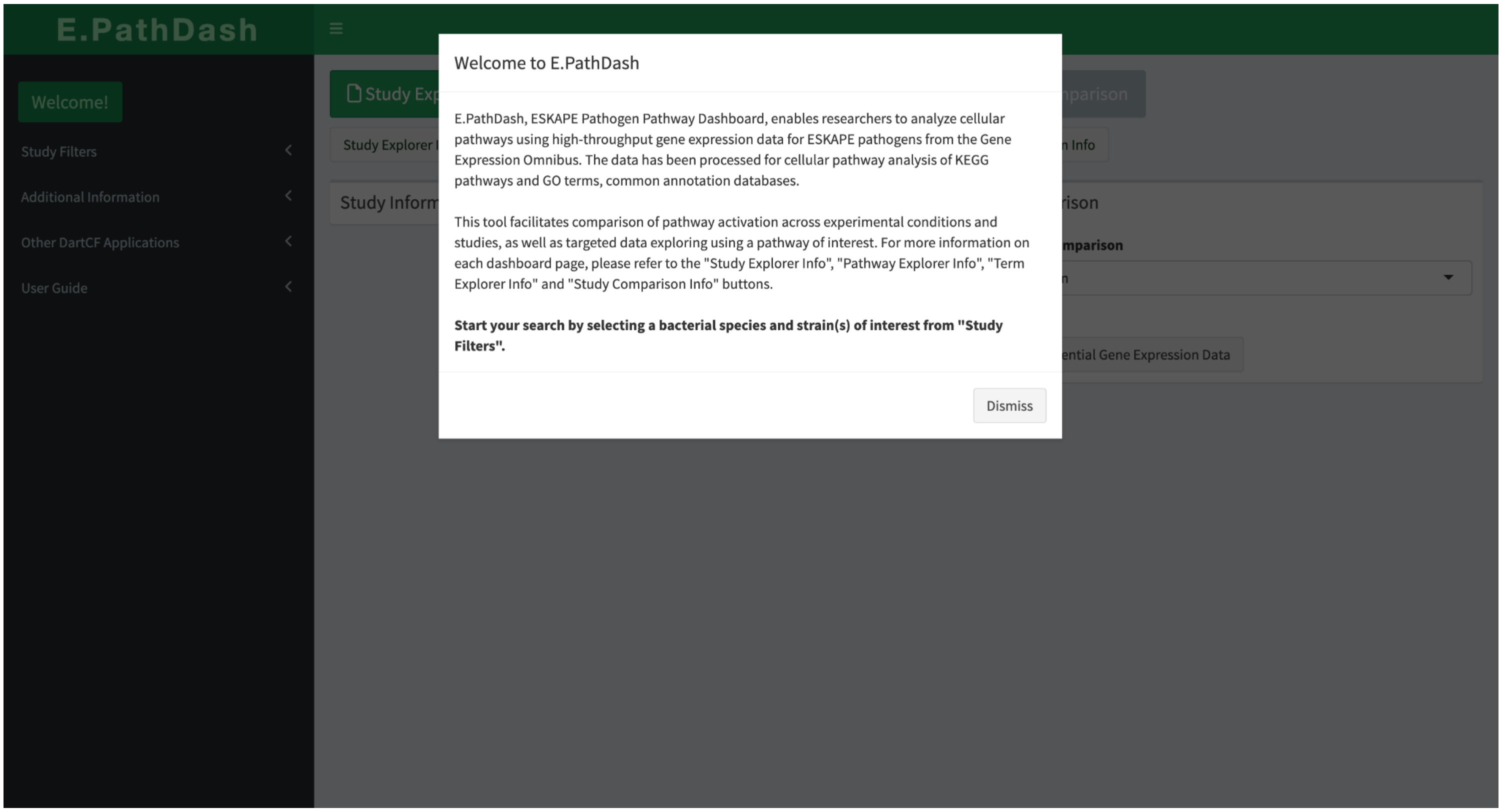

**Figure S1. Landing page**

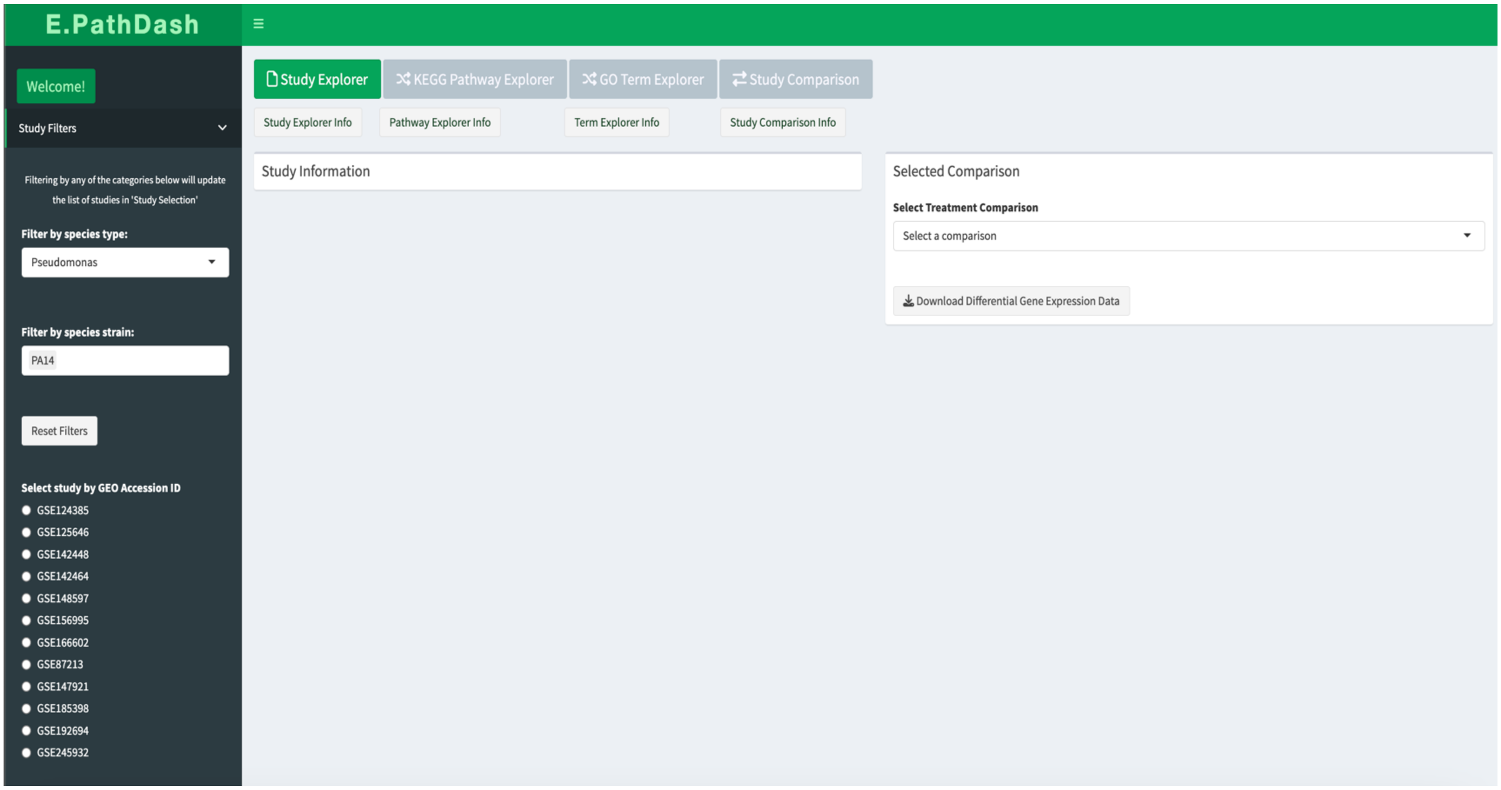

**Figure S2. Filtering side panel**

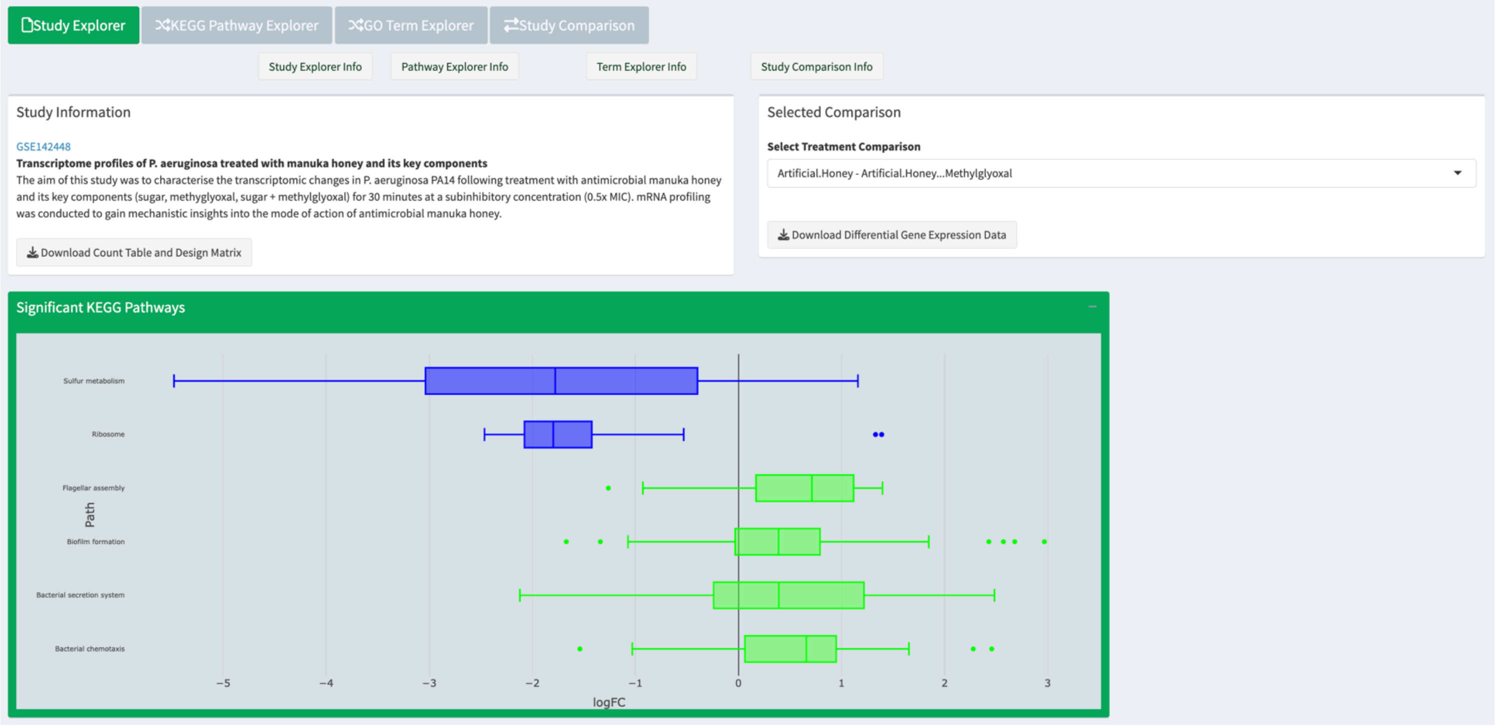

**Figure S3. Study Explorer page (section 1)**

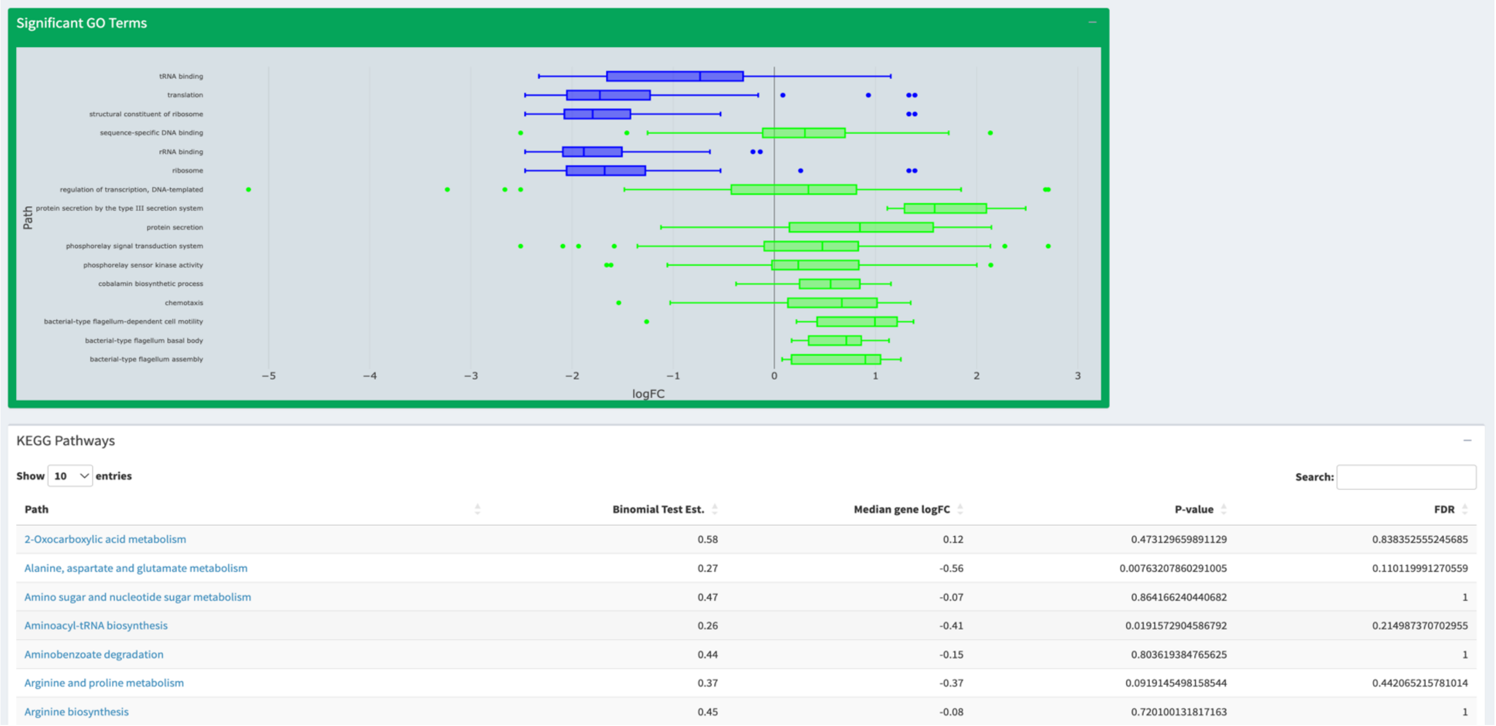

**Figure S4. Study Explorer Page (section 2)**

| | | | | |
|---|---|---|---|---|
| Aminobenzoate degradation | 0.44 | -0.15 | 0.803619384765625 | 1 |
| Arginine and proline metabolism | 0.37 | -0.37 | 0.0919145498158544 | 0.442065215781014 |
| Arginine biosynthesis | 0.45 | -0.08 | 0.720100131817163 | 1 |
| Ascorbate and aldarate metabolism | 0.43 | -0.06 | 1 | 1 |
| Bacterial chemotaxis | 0.79 | 0.66 | 0.0000616964077764238 | 0.00124626743708376 |
| Bacterial secretion system | 0.67 | 0.39 | 0.000924644411771929 | 0.0155648475981608 |

Showing 1 to 10 of 101 entries

Previous 1 2 3 4 5 … 11 Next

Download Table

#### GO Terms

Show 10 entries

Search:

| GO Term | Binomial Test Est. | Median gene logFC | P-value | FDR |
|---|---|---|---|---|
| 'de novo' IMP biosynthetic process | 0.14 | -0.81 | 0.012939453125 | 0.186697823660714 |
| 'de novo' UMP biosynthetic process | 0.29 | -0.61 | 0.453125 | 0.888470852759577 |
| 2 iron, 2 sulfur cluster binding | 0.45 | -0.02 | 0.720100131817163 | 1 |
| 3-deoxy-7-phosphoheptulonate synthase activity | 0.4 | -0.55 | 1 | 1 |
| 3-oxoacyl-[acyl-carrier-protein] synthase activity | 0.43 | -0.39 | 1 | 1 |
| 3'-5' exonuclease activity | 0.25 | -0.42 | 0.2890625 | 0.695126488095238 |
| 4 iron, 4 sulfur cluster binding | 0.36 | -0.24 | 0.0396170148984998 | 0.300098887856136 |
| ABC-type amino acid transporter activity | 0.83 | 0.19 | 0.21875 | 0.607508296460177 |
| ABC-type sulfate transporter activity | 0 | -3.2 | 0.0625 | 0.32876786023016 |
| acetyl-CoA carboxylase activity | 0.5 | -0.07 | 1 | 1 |

Showing 1 to 10 of 303 entries

Previous 1 2 3 4 5 … 31 Next

Download Table

**Figure S5. Study Explorer page (section 3)**

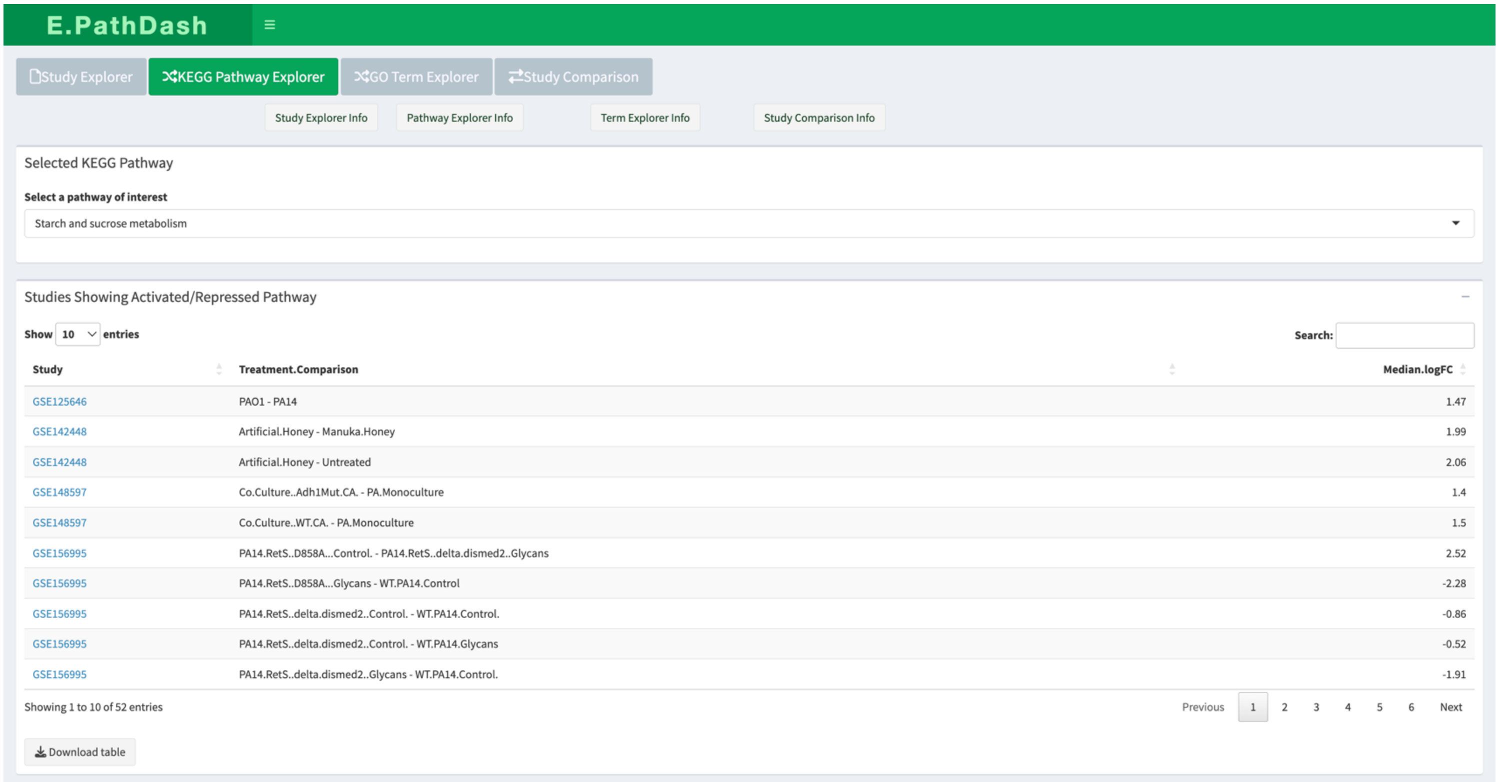

**Figure S6. KEGG Pathway Explorer page (section 1)**

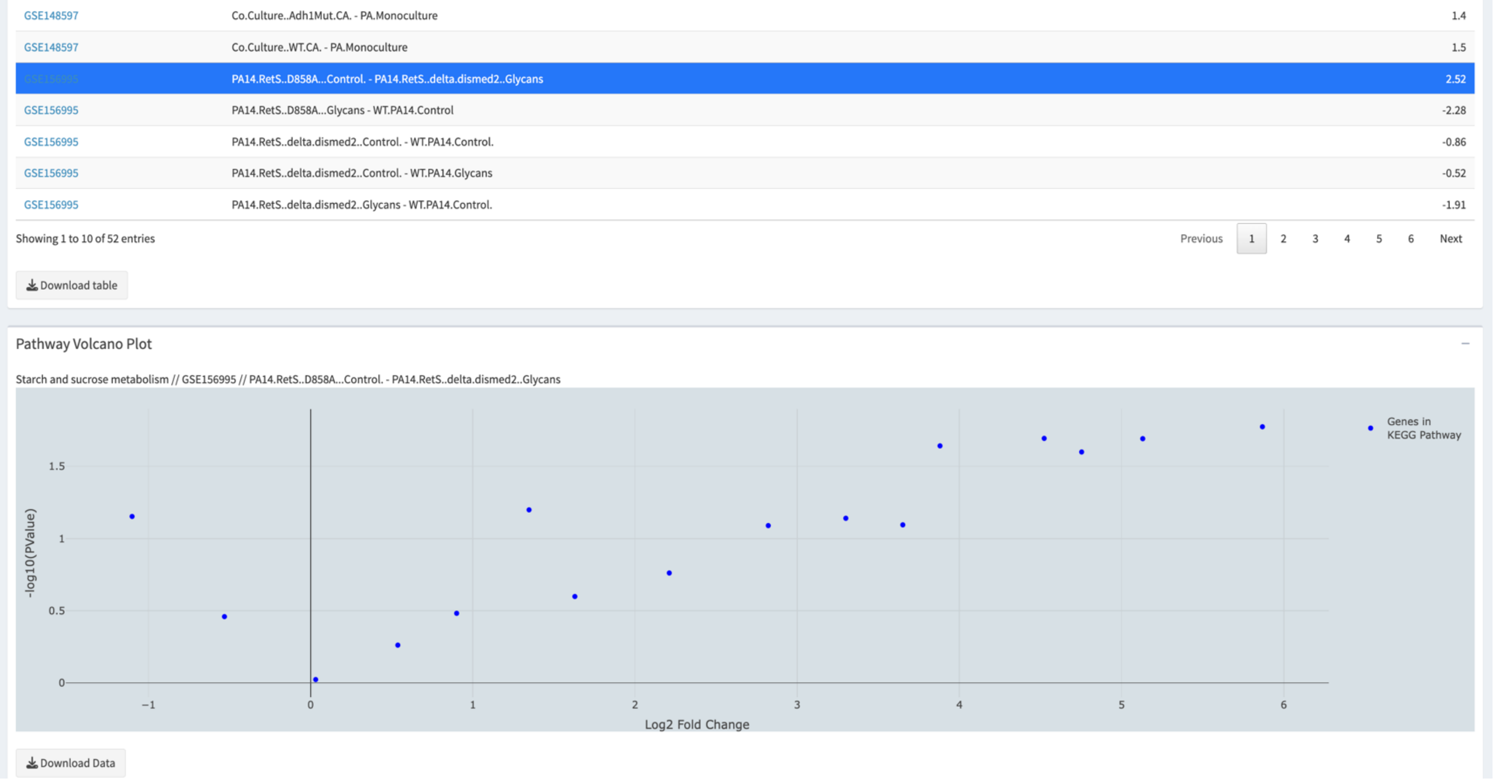

**Figure S7. KEGG Pathway Explorer (section 2)**

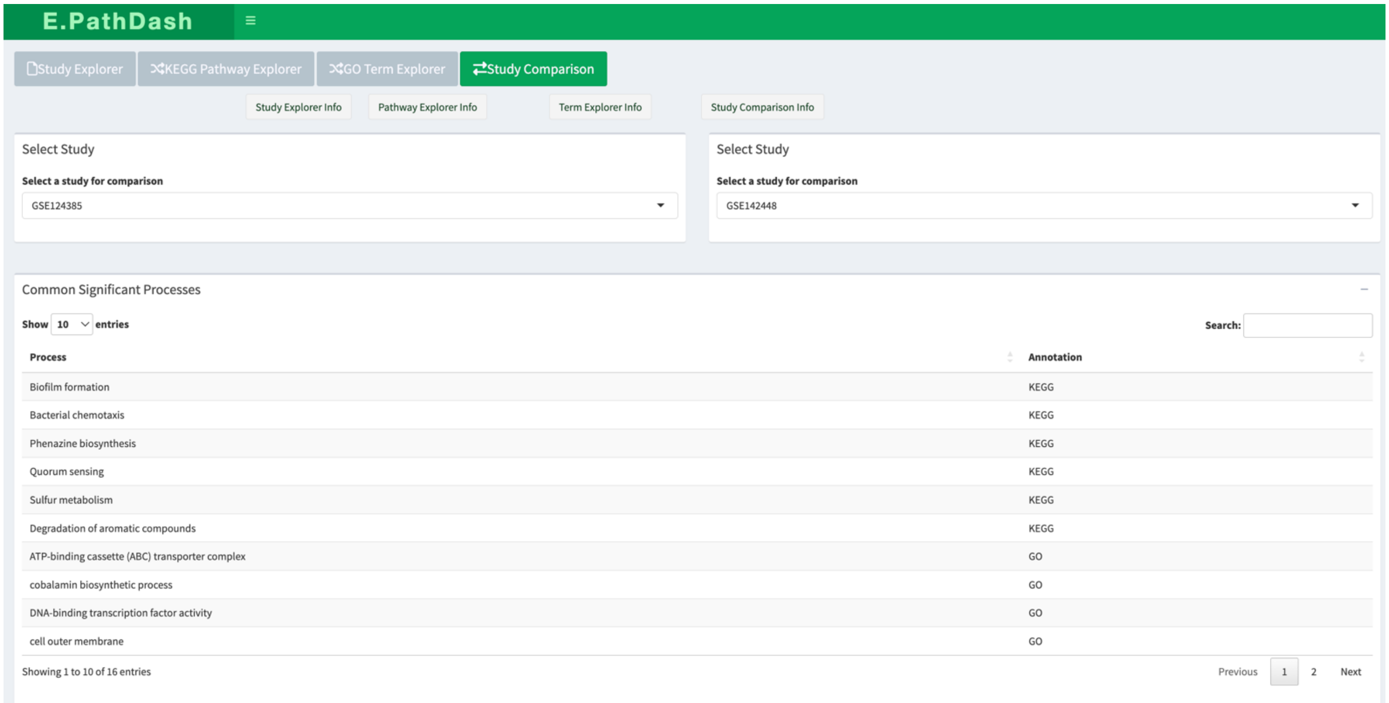

**Figure S8. Study Comparison page (section 1)**

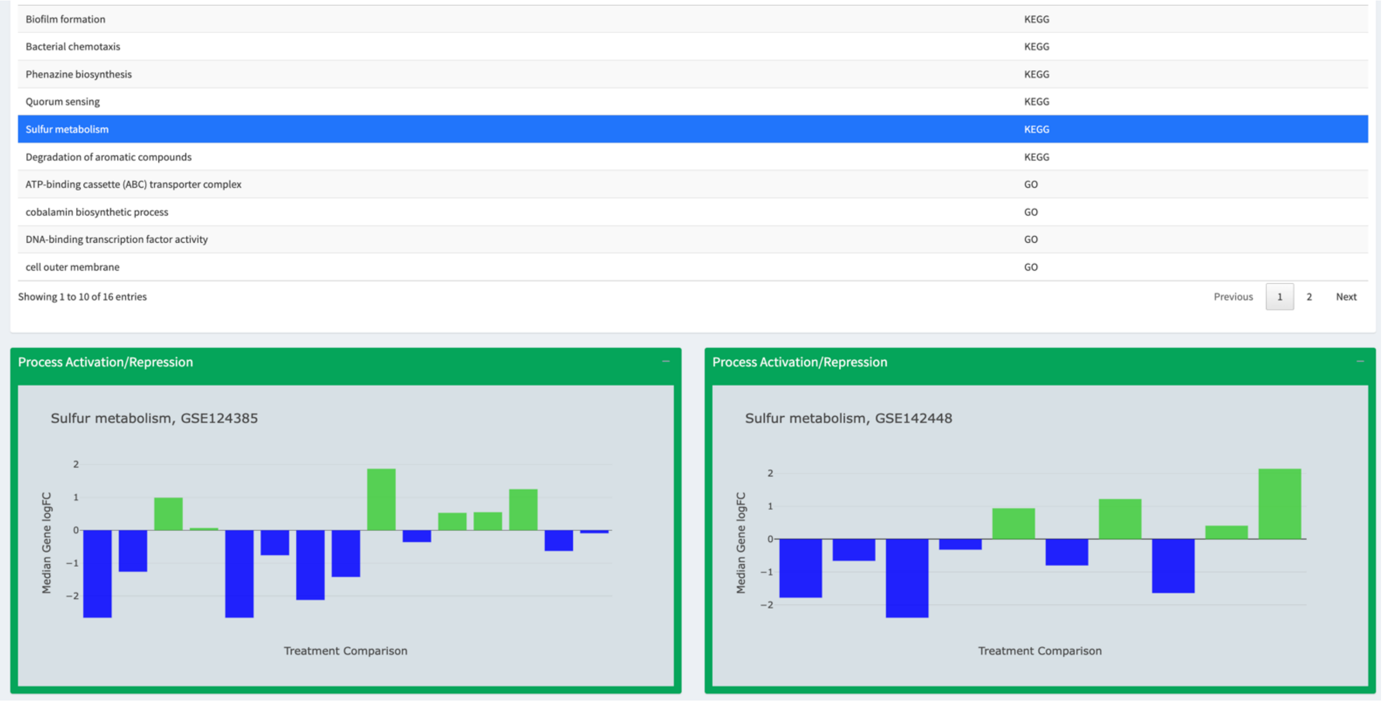

**Figure S9. Study Comparison page (section 2)**

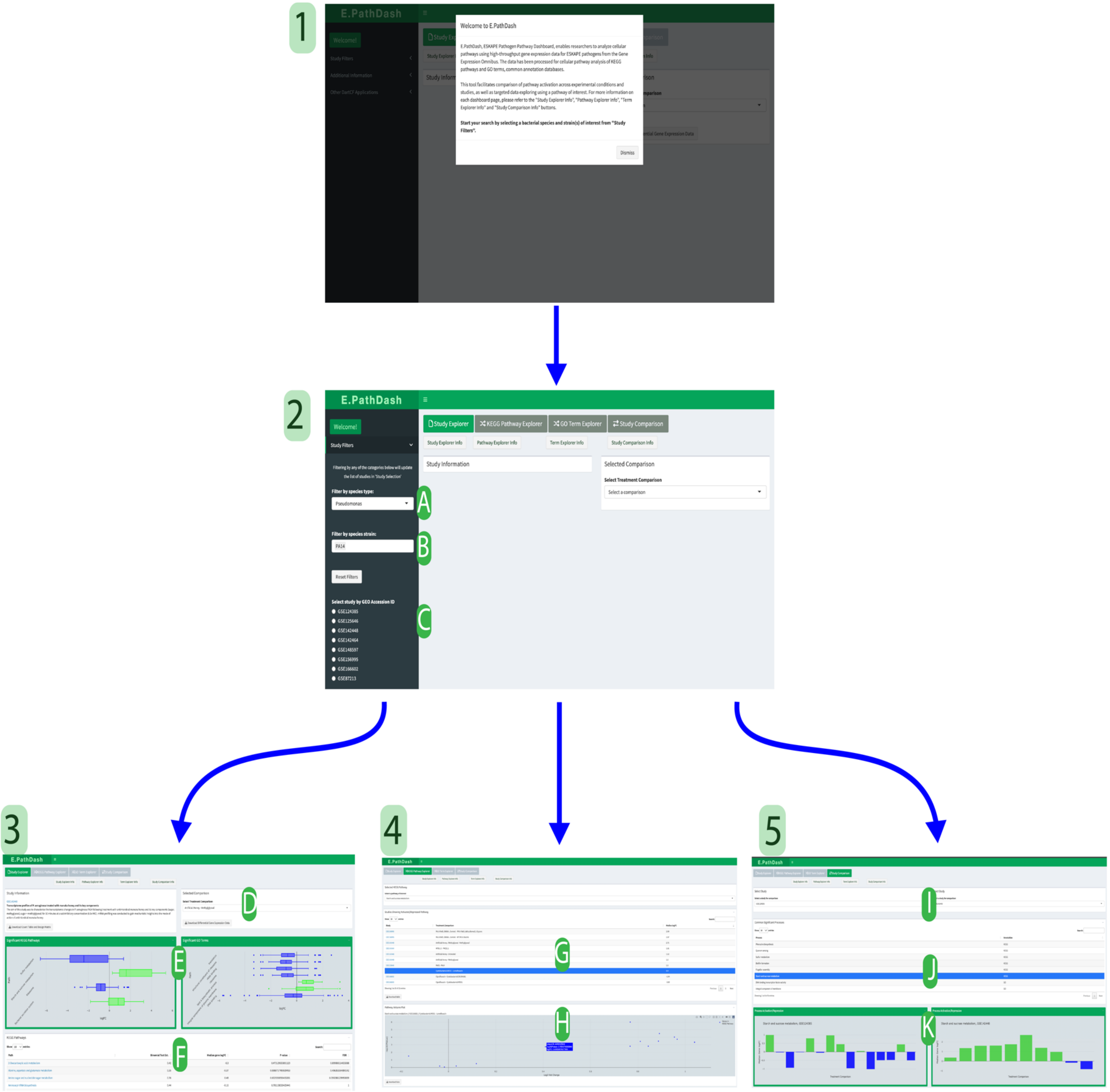

**Figure S10. Enlarged Figure 2 from paper**
